# Immune signatures of common exposures through co-occurrence of T-cell receptors in tens of thousands of donors

**DOI:** 10.1101/2024.03.26.583354

**Authors:** Damon H. May, Steven Woodhouse, Matthew T. Noakes, Kathryn Doroschak, Rebecca Elyanow, Amanda J. Moore, Edward J. Osborne, Brad Greenfield, Tim Hayes, Ruth Taniguchi, Zheng Yang, John R. Grino, Rachel Byron, Jamie Oaks, Haiyin Chen-Harris, Anna Sherwood, Julia Greissl, Bryan Howie, Mark Klinger, Harlan S. Robins

## Abstract

**Background:** Memory T cells are records of clonal expansion from prior immune exposures such as infections, vaccines and chronic diseases. Some of the receptors of these expanded T cell clones in a typical immune repertoire are highly public (present in many individuals) because they respond to the same peptide from a prevalent immune exposure, presented by the same Human Leukocyte Antigen (HLA) allele. Only a tiny fraction of public T-cell receptor β sequences (TCRs) have known associations with exposures or specific peptides.

**Methods:** We mined the TCR repertoires of tens of thousands of donors to define “ECOclusters”: clusters of public TCRs that tend to occur in the same donors. First, we built models to infer donor HLA type from the TCR repertoire, then associated public TCRs with HLA alleles. Next, we derived co-occurrence clusters of TCRs responding to antigens presented by the same HLA allele, then combined those clusters by co-occurrence across HLA alleles. Each such cross-HLA ECOcluster putatively represents a public TCR signature of a single exposure.

**Results:** We constructed sensitive, specific models to predict the presence of 220 HLA alleles from TCR repertoires and clustered 8,618,285 HLA allele-associated TCRs to define 11,058 ECOclusters. Using serologically labeled repertoires, we identified ECOclusters associated with HSV-1, HSV-2, EBV, Parvovirus, *Toxoplasma gondii*, Cytomegalovirus and SARS-CoV-2, and constructed sensitive, specific classifiers of exposure. ECOclusters represent a step toward deciphering the ledger of immune exposure history encoded by the T-cell repertoire.

## 1 Introduction

The enormous diversity of T cells in any individual allows for immune system recognition of many foreign pathogenic exposures. A given individual’s T-cell repertoire is a mix of naive and memory T cells, largely shaped by the combination of naive T-cell generation early in life and the exposure history of the individual (1–3).

T cells are activated when T-cell receptors recognize cognate antigens (4) presented by major histocompatibility complexes known as human leukocyte antigen (HLA) molecules in humans. HLA genes are the most polymorphic in the genome. Different HLA alleles (defined here with 4-digit precision, e.g., A*02:01) present complementary sets of antigens, and T-cell receptors are often observed to be specific to a combination of peptide antigen and restricting HLA allele (pHLA) (5–7).

T cells in subjects sharing both an HLA allele and a common immune exposure will encounter some of the same pHLAs.

Since this work deals with bulk TCRβ sequencing, we define a TCR as a productive combination of TCRβ V gene, J gene and CDR3 amino acid sequence. High-throughput sequencing of a peripheral blood sample may measure ∼10^6^ TCRs (8). An individual’s immune history is encoded in these repertoire TCRs (9). However, given the generally unknown pHLA and peptide associations of TCRs, the high-dimensional nature of repertoires and the genetic diversity of individuals expressed in their inherited HLA types, disentangling the many signals present in a repertoire presents many challenges (10–12).

Subjects who share a prior immune exposure tend to share some TCRs responding to the exposure. It has been previously shown that these “public” TCRs can be associated with immune exposures and used to build diagnostic models for exposures such as Cytomegalovirus (CMV) (13), SARS-CoV-2 (14), Lyme disease (15), herpes simplex virus 1 and 2 (HSV-1/HSV-2) (12) and others. The exposure-associated TCRs bind antigens derived from the exposure, each within the context of an HLA allele restriction (Figure 1A).

**Figure 1.**
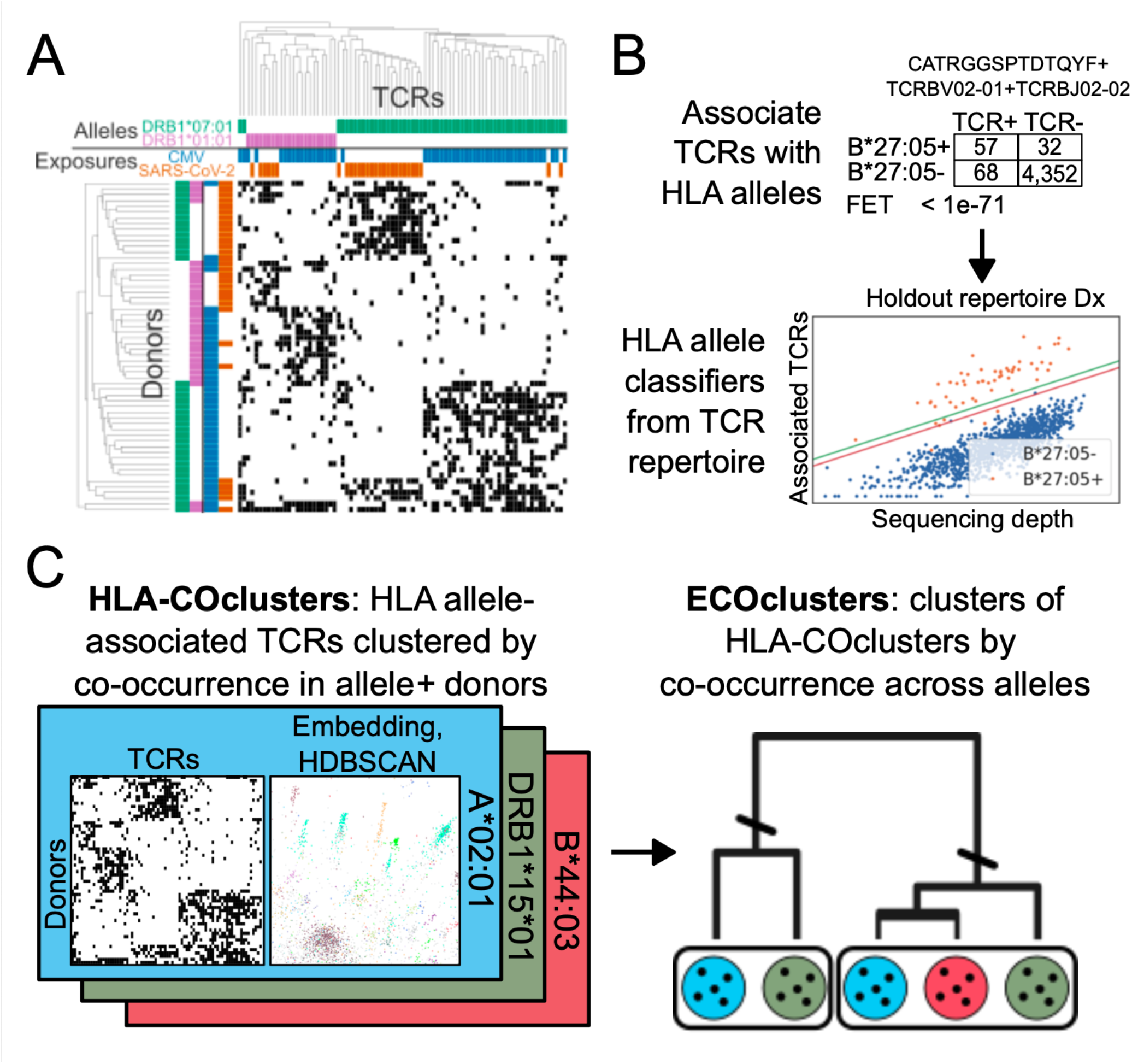
A. Illustration of TCR clustering by repertoire HLA type and exposure history. Heatmap shows presence (black) or absence (white) of 80 TCRs in 58 donor repertoires. TCRs have known HLA allele association (green or pink) and association with exposure to CMV (blue) or SARS-CoV-2 (orange). Donors have known positive (color) or negative (white) HLA alleles and exposures. TCRs and repertoires cluster (here, average-linkage hierarchical clustering) primarily by HLA allele and secondarily by exposure status. B. Building HLA classifiers. We associate TCRs with HLA alleles by occurrence in allele+ and allele-donors and train a predictive model predicting HLA allele presence (orange) or absence (blue) on allele-associated TCR count and sequencing depth, defining positive (green) and negative (red) call thresholds. C. Constructing ECOclusters. Separately for each HLA allele, cluster HLA allele-associated TCRs by co-occurrence within allele+ donors. Cluster these clusters by donor occurrence correlation, considering only donors having the allele(s) associated with both clusters.

Similarly, by comparing TCR repertoires from subjects expressing *vs*. not expressing a particular HLA allele, we and others have associated public TCRs with HLA alleles (16–18). Each such TCR is associated with a single HLA allele due to its response to an HLA-presented antigen from some prevalent exposure.

Here, expanding on ideas first articulated by DeWitt et al. (9), we introduce an approach to identify public TCRs responding to prevalent exposures using the co-occurrence patterns of TCRs observed in tens of thousands of TCR repertoires. We first associate public TCRs with the HLA alleles that present the antigens they bind and use those TCRs to build repertoire-based classifiers for HLA allele presence (Figure 1B). Next, we use those classifiers to infer donor HLA type from 34,793 repertoires and then build a much larger database of HLA allele-associated TCRs. We then construct clusters of co-occurring public TCRs, in two stages (Figure 1C): separately for each HLA allele, we identify HLA-COclusters (HLA Co-Occurrence clusters), clusters of TCRs associated with the same HLA allele that co-occur in repertoires from donors inferred to have the allele; then, we cluster the HLA-COclusters by co-occurrence across HLA allele associations to derive ECOclusters (Exposure Co-Occurrence clusters). Each ECOcluster putatively represents a public TCR response to a single prevalent exposure.

We validate this approach using repertoires with serological labels for seven prevalent exposures. For each exposure, we identify associated ECOclusters that discriminated serological cases from controls in a held out set of labeled repertoires. We also “deorphanize” several ECOclusters through their high concentration of TCRs known to bind to antigens presented by the same exposure, decoding more of the public T-cell repertoire.

An earlier version of ECOclusters was described in two 2024 preprints (19,20). Since that time, further published papers and preprints have confirmed the clinical relevance of ECOclusters to Cytomegalovirus (21), Primary Sclerosing Cholangitis (22) and Celiac disease (23). The work described here represents a significant expansion on the earlier ECOclusters preprints. Here, we describe ECOclusters built from repertoires from more-diverse donors, incorporating TCRs associated with many more HLA alleles and associating ECOclusters with more exposures.

## 2 Materials and Methods

### 2.1 Sample acquisition and TCR repertoire sequencing

We used previously sequenced TCR repertoires from 4,144 primarily self-reported White and Black donors of known HLA type described previously (24). To increase the diversity of HLA alleles represented, we acquired samples from 1,328 donors with self-reported Asian and other ancestry in the USA and from 482 Kuwaiti donors (Supplementary Figure 1). HLA genotyping was obtained from Scisco Genetics using the ScisGo-HLA-v6 methodology. We performed immunosequencing as previously described (8,14), bringing our total to 5,954 HLA-typed repertoires.

Between previously sequenced and newly sequenced repertoires, we obtained TCR repertoires for 46,080 donors from 13 cohorts (summary in Supplementary Table 1). The 5^th^-95^th^ percentile range of the total number of unique rearrangements sequenced per donor was 93,278-562,172, with median 305,253.

### 2.2 Constructing classifiers for HLA allele status

We built classification models for each of 220 HLA alleles (where an “HLA allele” can be an individual allele, a class II DP/DQ heterodimer or a p group – a set of HLA alleles with identical antigen binding domain protein sequences (25)) with a modified version of the methodology previously described (24) (Supplementary Figure 1C, details in Supplementary Methods).

These 220 models differ from those described in Zahid et al. in including more training data from more diverse donors, modeling more HLA alleles and introducing separate positive and negative call thresholds (and allowing models to abstain on some repertoires). We also modeled some sets of HLA alleles collectively as p groups, after observing that models trained individually were essentially indistinguishable in our data or that high-precision models for alleles within a p group could not be trained individually.

### 2.3 Associating public TCRs with HLA alleles

To construct a database of ^∼^8.6 million HLA allele-associated public TCRs, we used our HLA classifiers to computationally HLA type a cohort of 34,793 donors with respect to the 220 modeled HLA alleles. We then identified candidate TCRs overrepresented in predicted positives for each HLA allele with Fisher’s Exact Test and estimated a False Discovery Rate (FDR, method described in Supplementary Methods) for each TCR. We retained TCRs with *p* < 0.0001 and FDR < 0.001. For TCRs with multiple retained allele associations, we chose the allele with the lowest FDR.

### 2.4 Clustering TCRs by co-occurrence in repertoires

For each HLA allele *a*, we then constructed a matrix indicating convergent recombination count of each *a*-associated TCR (the number of unique nucleotide rearrangements encoding the TCR) in each repertoire imputed to have *a*. Clustering on these matrices proved to be intractable due to their extreme sparsity. We therefore first transformed each matrix through a series of embedding and dimensionality reduction steps.

For each such matrix, we embedded both TCRs and donors into a shared space of 150 dimensions using spectral co-clustering (26). The input bipartite graph is between TCRs and donors, and edges are weighted by convergent recombination count. This embedding reduces the dimensionality and sparsity of the data and relates the problem of clustering TCRs in terms of donors to the problem of clustering donors in terms of TCRs. Next, we applied UMAP (27) to further reduce the dimensionality to 15, which enabled application of the density-based clustering algorithm HDBSCAN (28) with a minimum cluster size of 10 TCRs and/or donors. HLA-COclusters are the TCR members of each derived cluster.

Next, we built cross-HLA ECOclusters by clustering HLA-COclusters across allele associations. Let *X* denote the matrix of donors by HLA-COclusters, with values as summed convergent recombination counts across the HLA-COcluster TCRs. We defined an HLA-masked correlation matrix *C* on all pairs of HLA-COclusters:

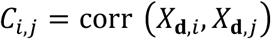

where corr denotes Pearson correlation and **d** denotes the donors imputed to have the HLA allele or alleles associated with both HLA-COclusters *i* and *j*.

We then performed agglomerative hierarchical clustering using 1 – *C* as a pairwise distance between HLA-COclusters and a custom linkage function *f*:

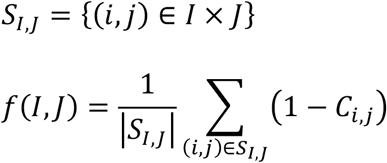

*f* preserves the average HLA-masked correlation distances between sets of HLA-COclusters at every branch point with higher fidelity than standard linkage functions such as the UPGMA linkage, which is defined by repeated averaging.

Finally, we cut the resulting hierarchical clustering tree to define a discrete set of clusters, using natural cut points learned from the structure of the tree. We observed that most paths from leaf HLA-COclusters to the root of the tree exhibited sudden spikes in the value of *f* at which many HLA-COclusters come together. Accordingly, for each leaf HLA-COcluster, we located the branch point maximizing the second derivative of *f, f’’*, along the path from root to leaf. Many HLA-COclusters share a maximum *f’’* point, converging to the same cluster assignment. However, along some branches we obtained nested clusters. In these cases, we retained the cluster closest to the root.

### 2.5 Quantifying ECOcluster breadth in repertoires

To quantitatively measure the TCR burden of an ECOcluster *e* within a repertoire *r*, we defined an HLA-invariant estimate of the “breadth” of *e* within *r, B*_*er*,_. An ECOcluster *e* comprises HLA-COclusters associated with a set of HLA alleles *A*_*e*_. A repertoire *r* has a set of inferred HLA alleles *A*_*r*_. Define *A*_*re*_ as the intersection of *A*_*r*_ and *A*_*e*_. Each repertoire can have a different *A*_*re*_ with respect to the same ECOcluster *e*; therefore, *B*_+,_ is undefined if *A*_*re*_ is the empty set, and it must account for *A*_*re*_ in order to be comparable across repertoires.

Conceptually, we build *B*_*er*_ on ECOcluster *e* within repertoire *r* as follows: calculate the simple breadth of each HLA-COcluster associated with an HLA allele in *A*_*re*_; normalize the breadth on each HLA-COcluster *h* as its percentile within the breadths on *h* from a large reference distribution of repertoires; calculate *B*_*er*_ as the weighted mean of those breadths (using different weights in different contexts, see below). *B*_*er*_ is interpretable as an estimate of *r*’s percentile within a notional HLA-independent distribution of breadths on *e*. A formal definition follows.

Define *N*_*r*_ as the total number of unique rearrangements observed in *r*, and *C*_*hr*_ as the number of rearrangements in both repertoire *r* and HLA-COcluster *h*. The simple breadth of HLA-COcluster *h* in repertoire *r, B*_*hr*_, is calculated as:

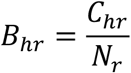

We wish to compare breadth values *B*_*hr*_ within the same ECOcluster between donors having different HLA types and therefore having nonmissing *B*_*hr*_ values for different HLA-COclusters. However, HLA-COclusters differ in repertoire impact, due to e.g. varying TCR count and cognate peptide immunogenicity, and *B*_*hr*_ values for different HLA-COclusters are not directly comparable. We therefore normalize *B*_*hr*_ by calculating 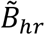, the percentile of *B*_*hr*_ within all the breadth values for *h* from a large database of repertoires (the repertoires used in the construction of ECOclusters) inferred to have the HLA allele *a* associated with *h*. That is:

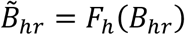

where *F*_*h*_ is the empirical cumulative distribution function on *B*_-,_for repertoires inferred to have *a*. 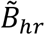, then, has the same interpretation for each HLA-COcluster within the same ECOcluster: a percentile-transformed measure of each donor’s response to the exposure represented by the ECOcluster, among all HLA-matched donors.

Finally, some HLA-COclusters may be thought to be more salient or more accurately measured than others (see below), so their 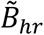, values should be weighted more highly. Let w_*h*_ denote some nonnegative weight assigned to HLA-COcluster *h*. We define a measure of the breadth of ECOcluster *e* in repertoire *r, B*_*er*_, as the weighted mean of the normalized breadths of the HLA-COclusters in *e* HLA-matched to *r*. More formally, for repertoire *r* with inferred HLA alleles A_*r*_, define the set of HLA-COclusters *H*_*r*_ = {*h*:*h* is associated with an allele in *A*_*r*_}, and define *H*_*er*_ as *H*_*e*_ ∩ *H*_*r*_. Given weights *w*_*h*_ on each HLA-COcluster *h*, we calculate *B*_*er*_ as:

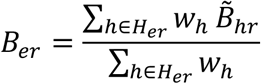

Intuitively and empirically, larger HLA-COclusters tend to have less dropout (repertoires containing no member TCRs) and therefore represent a more accurate measure of breadth. Therefore, as a general-purpose breadth measure, we define the weight of each HLA-COcluster *h* as *n*_*h*_, the count of TCRs in *h*, to derive 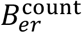:

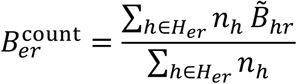

An alternative weighting scheme specific to the context of groups of serologically labeled repertoires is described below.

### 2.6 Constructing sets of serologically labeled repertoires

For seven prevalent exposures, we assembled collections of repertoires from donors with positive or negative serolabels for each exposure. For CMV and SARS-CoV-2, we used previously acquired serologically labeled samples as described previously (13,14). For EBV, Parvovirus, HSV-1, HSV-2 and *T. gondii*, we derived new serological labels on previously acquired samples using U-PLEX Development Pack from Meso Scale Discovery (see Supplementary Methods for details).

For each indication, labeled repertoires were divided into training and holdout sets. Where possible, this division separated meaningfully different cohorts. The holdout set samples for EBV and *T. gondii* were acquired with the explicit aim of increasing diversity with respect to HLA type and race/ethnicity (see above). All holdout repertoires from all indications were excluded from the construction of the HLA allele-associated TCR database and ECOclusters.

The samples labeled for HSV-2 were from a single cohort, but with a mix of self-reported Black and White donors. We split the labeled repertoires into a training set of White donors and a holdout set of all Black donors. Separately within the training and holdout sets, cases and controls were matched on age and sequencing depth. These training and holdout sets were provided to support AIRR-ML-25, a completed public machine learning competition (https://www.kaggle.com/competitions/adaptive-immune-profiling-challenge-2025) whose results will be published in Nature Methods.

### 2.7 Associating ECOclusters with serological labels

For each indication, we calculated repertoire breadth 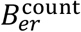 as described above for every ECOcluster on all repertoires in the training set. We tested each ECOcluster for differential 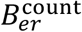 in cases using a two-sided Mann-Whitney U test (MWU) on serologically positive *vs*. negative 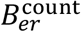 for each ECOcluster, applying Bonferroni correction for multiple testing. For all ECOclusters with Bonferroni-corrected MWU *p* < 0.001, we calculated sensitivity at 98% specificity for 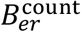 as a predictor of case status within the training set. We considered all ECOclusters with higher mean 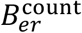 in cases than controls, MWU *p* < 0.001 and >20% sensitivity at 98% specificity to be associated with the indication.

### 2.8 Constructing classifiers for prior immune exposure

For each indication, for each indication-associated ECOcluster *e*, for each HLA-COcluster *h* within *e*, we calculated the ROC curve relating *B*_*hr*_ to the positive serological label in the training set. For our classifier, we retained all HLA-COclusters with area under the receiver-operator characteristic curve (AUROC) >0.6, >20% sensitivity at 98% specificity and at least 5 positive and 5 negative labels in HLA-matched repertoires (the 5-positive requirement was waived for *T. gondii* due to low prevalence in training samples).

Whether they came from a single ECOcluster or multiple, we treated all retained HLA-COclusters for an indication as though they were members of a single ECOcluster *e* and calculated an alternatively-weighted *B*_*er*_ breadth measure for this task, as described above, with one difference: the weights for each HLA-COcluster *h* in this breadth measure were defined by the AUROC for *h* in the training set:

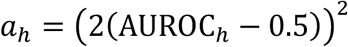

To classify new repertoires with respect to the label, we used a weighted breadth 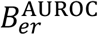 calculated using these weights *a*_*h*_:

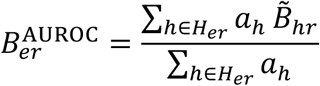

We chose the 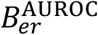 threshold for binary classification based on the training ROC curve, as the value maximizing Youden’s *J* statistic (*J* = true positive rate – false positive rate).

Alternative models (e.g., gradient-boosted trees) were considered that more directly inferred each HLA-COcluster’s contribution to the classification task. We chose this model because it outperformed those models in cross-validation and because 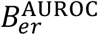 is interpretable as an HLA-independent estimate of the donor’s breadth on exposure-responding TCRs, expressed as a percentile in a large reference distribution.

### 2.9 Computing enrichment of ECOclusters for TCRs with known association

To associate ECOclusters with exposures for which we lacked labeled repertoires, we evaluated ECOclusters for statistically unlikely representation of TCRs with known association with the same taxon. We separately considered two sources for those associations: public databases and our MIRA assay (29), which identifies TCRs from T cells activated by specific peptides.

We combined three public databases of associations between TCRs and peptide antigens: VDJDB (score > 1) (30), IEDB (31) and McPAS(32), downloaded from their respective websites, into a single dataset. In this analysis, because of differences in gene naming between our data and the databases, we matched TCRs by CDR3 amino acid sequence, V gene family and J gene family. For the MIRA analysis, we used V gene, J gene and CDR3 amino acid sequence.

Given each of these datasets of TCR-pHLA associations, we separately tested each ECOcluster for enrichment of TCRs associated with pHLAs from each represented taxon using a method analogous to gene set enrichment analysis. For each combination of ECOcluster and taxon, we calculated the number *x* of unique TCRs shared between the ECOcluster and the taxon-associated TCR list. We then computed the probability *p(x)* of observing an intersection of *x* or more TCRs with an ECOcluster by chance, using the hypergeometric distribution:

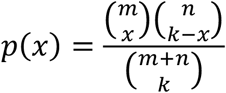

where:

*x* = count of intersecting TCRs

*m* = count of TCRs in ECOcluster

*k* = count of TCRs in TCR list

*n* = (estimate of total number of public TCRs) - *m*

For the total number of public TCRs *n + m*, smaller values are more conservative in the sense of leading to higher *p*-values for a given intersection. Therefore, we estimated this value conservatively as 8,618,285, the size of our HLA allele-associated TCR database and therefore the smallest possible estimate. We retained associations with *p* < 1e-10 and at least 15 intersecting TCRs associated with at least 2 different peptides.

## 3 Results

### 3.1 HLA-associated public TCRs from tens of thousands of repertoires with imputed HLA

Most public TCRs notionally respond to a cognate antigen from a prevalent immune exposure presented by the MHC encoded by an HLA allele or else have extremely high V(D)J generation probability (pGen, here computed by OLGA (33)). As described above, a public TCR’s pattern of occurrence across a group of donor repertoires tends to be defined by which donors have its corresponding HLA allele and which donors have encountered its associated exposure.

Accordingly, to discover groups of public TCRs associated with exposures, we first inferred donor HLA type with respect to 220 HLA alleles on 34,793 donors and associated 8,618,285 of their public TCRs with HLA alleles (see Supplementary Results for details). 92% of TCRs were associated with Class II HLA alleles and 8% with Class I alleles. We held out from this process all repertoires later used to evaluate diagnostic model performance.

### 3.2 Clusters of co-occurring TCRs in tens of thousands of repertoires

We next clustered HLA allele-associated TCRs by their co-occurrence in donor repertoires.

First, we built *HLA-COclusters*: TCRs that co-occur in donors within the context of an HLA allele. Separately for each HLA *h*, we embedded donors inferred positive for *h* and TCRs associated with *h* into a shared space using an embedding adopted from the spectral co-clustering algorithm (26) and then reduced this space to 15 dimensions by applying UMAP (27). We note that any method for dimensionality reduction can lose important features of the correlation structure of a dataset, but we found these steps necessary to deal with the extreme sparsity and high dimensionality of the data (the “curse of dimensionality” in machine learning (34). We used density-based clustering (28) to derive TCR clusters, obtaining 44,010 HLA-COclusters across 196 HLA alleles.

To bridge across HLA contexts, we built *ECOclusters*: clusters of HLA-COclusters that span one or more HLA alleles. To do this, we computed an HLA-masked correlation matrix between all pairs of HLA-COclusters whose entries are the Pearson correlation coefficient between pairs of HLA-COclusters, in only the donors that are positive for the HLA allele(s) associated with the pair. We then applied hierarchical clustering and cut the clustering tree at naturally defined “elbows” to derive 11,058 ECOclusters. ECOclusters contain between 6 and 207,512 TCRs, associated with between 1 and 81 HLA alleles (Figure 2A).

**Figure 2.**
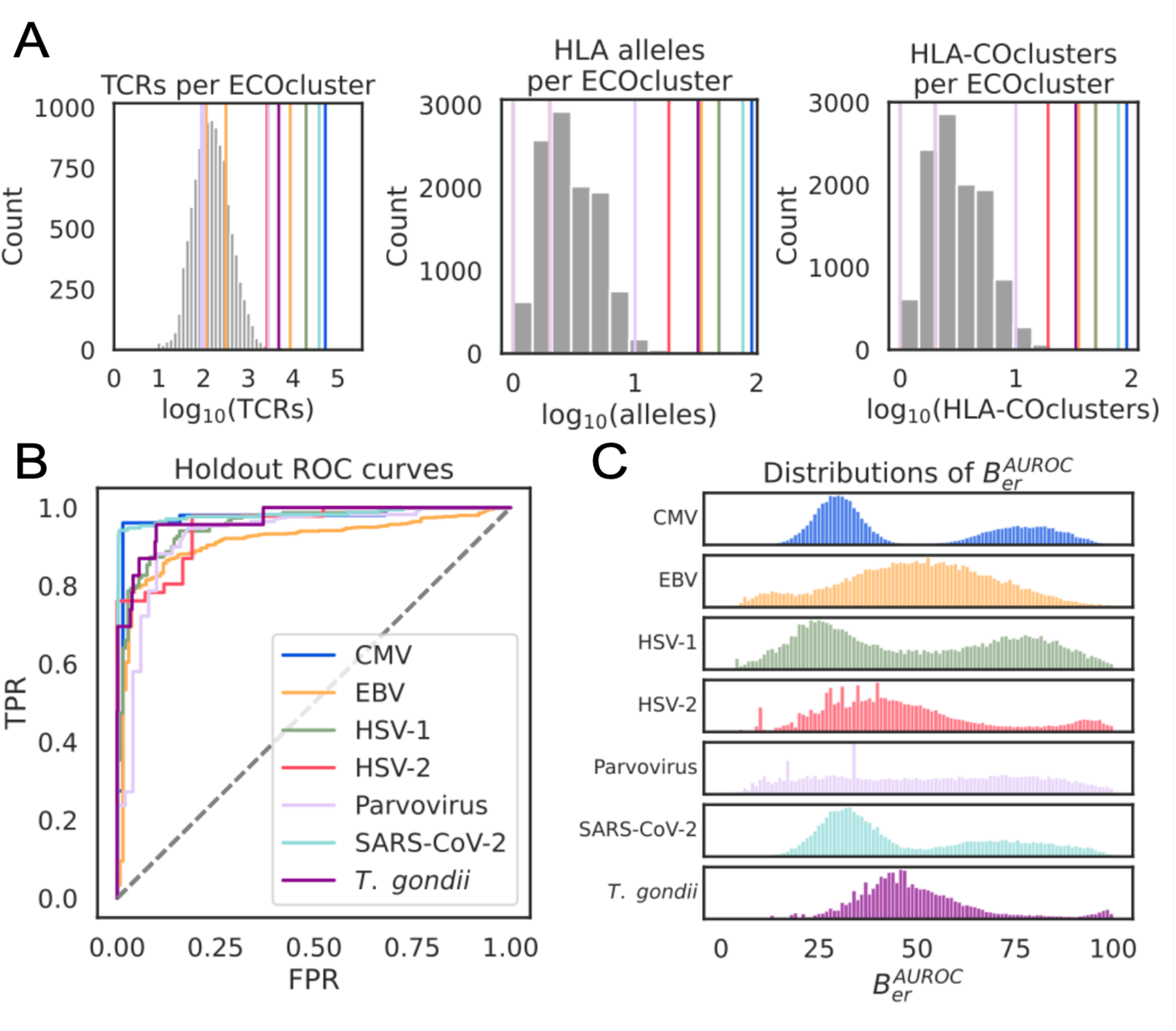
A. Histograms of the number of TCRs, HLA-COclusters, and unique HLA allele associations represented by each ECOcluster. ECOclusters used in diagnostic models indicated with colored lines. B. ROC curves describing the performance of 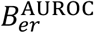 (estimated HLA-independent breadth) on case-associated ECOclusters as a classifier for seven prevalent exposures on held out labeled repertoires. C. Distributions of 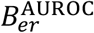 for ECOcluster associated with seven prevalent exposures within all donors from the 32,276-repertoire T-DETECT cohort inferred to have at least one HLA allele contributing to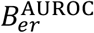. Distributions exhibit bimodality ranging from weak (Parvovirus) to nearly complete (CMV) separation between high (presumed exposed) and low (presumed unexposed) modes.

### 3.3 ECOclusters associated with specific exposures

#### 3.3.1 Diagnostic models of infection from serologically labeled repertoires

For each indication (CMV, EBV, HSV-1, HSV-2, Parvovirus, SARS-CoV-2 and *Toxoplasma gondii*) we acquired repertoires from serologically labeled samples as described in Methods. Depending on labeled sample availability for each indication, we separated repertoires into train and holdout sets by cohort (samples acquired at different times and places), by self-reported race, or, when biologically meaningful separation was unavailable, by sequencing batch.

For EBV and *T. gondii*, holdout samples were acquired specifically to represent greater racial and HLA diversity than the training sets; EBV holdout samples were propensity matched on age, and *T. gondii* on unique productive rearrangements, due to observed imbalances. For HSV-1, HSV-2 and Parvovirus, training and holdout samples were acquired as part of the same cohort and then serologically labeled, minimizing confounding variables that differentiate positive and negative labels. Table 1 gives counts of training and holdout positive- and negative-labeled repertoires per indication (more details in Supplementary Table 1, Supplementary Figure 3). All holdout labeled repertoires for all indications were excluded when constructing our HLA allele-associated TCR database and ECOclusters.

**Table 1.** Classifier summary. For each indication, indicates how training and holdout data were separated; the number of positive and negative labels used in training and holdout; the number of ECOclusters, HLA-COclusters and TCR used; the percent of holdout samples on which model abstained; and the AUROC and sensitivity at 98% specificity of the classifier on holdout labeled repertoires.

| Indication | Train-holdout separation | +/- Training labels | +/- Holdout labels | ECOclusters | HLA-COclusters | TCRs | Percent abstention | AUROC | Sensitivity at 98% specificity |
| --- | --- | --- | --- | --- | --- | --- | --- | --- | --- |
| CMV | cohort | 289/352 | 51/69 | 1 | 79 | 50,217 | 0% | 0.97 | 96% |
| EBV | cohort, race | 1,434/72 | 750/150 | 4 | 28 | 13,437 | 7% | 0.91 | 53% |
| HSV-1 | batch | 521/414 | 167/155 | 1 | 25 | 9,914 | 8% | 0.95 | 64% |
| HSV-2 | race | 63/315 | 50/50 | 1 | 14 | 2,133 | 12% | 0.95 | 76% |
| Parvovirus | batch | 652/176 | 199/63 | 3 | 12 | 2,933 | 16% | 0.93 | 27% |
| SARS-CoV-2 | cohort | 1,130/2,669 | 172/3,845 | 1 | 65 | 37,704 | 0% | 0.98 | 94% |
| *T. gondii* | cohort, race | 17/1,345 | 25/750 | 1 | 17 | 3,644 | 5% | 0.97 | 69% |

In the training samples for each indication, for each repertoire *r*, for each ECOcluster *e*, we calculated 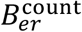, a general-purpose estimate of the HLA-independent contribution of *e* to *r* as a percentile of a reference distribution with respect to *e* (see Methods). We identified between 1 and 4 ECOclusters specifically associated with the positive label by 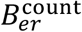 (four for EBV, three for Parvovirus and one each for the remaining labels), comprising between 12 and 81 HLA-COclusters and containing between 2,133 and 50,457 TCRs. For each label, we used the training set to learn positive-label-association weights for each associated HLA-COcluster and calculate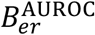: an estimate of HLA-independent contribution of the contribution of the associated HLA-COclusters to each repertoire that takes into account HLA-COcluster sensitivity and specificity and to set a classification threshold, defining a classifier (see Methods for details).

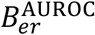is a more complex statistic than simple TCR breadth or HLA-aware breadth (i.e., matching TCRs in repertoires only if the donor is inferred to have the associated HLA allele), or even 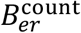.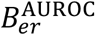 learns the diagnostic strength of each HLA-COcluster and then uses all HLA-COclusters that are matched to each repertoire, weighted appropriately. We found this additional complexity instrumental to extracting the highest-specificity diagnostic signal, particularly in indications that were more difficult to classify (Supplementary Figure 4 compares different breadth measures). As described previously (12), HSV-1 and HSV-2 share many common responding TCRs due to high protein homology, but the weights learned in constructing 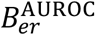 allowed those models to use the most indication-specific HLA-COclusters available to each donor.

#### 3.3.2 Performance of diagnostic models

In holdout labeled repertoires, our 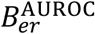-based classifiers achieved AUROCs ranging from 0.93 (Parvovirus) to 0.97 (CMV and SARS-CoV-2) and sensitivity at 98% specificity ranging from 27% (Parvovirus) to 96% (CMV). They abstained from prediction on any repertoires not inferred to have any of the HLA alleles used by the classifiers, with abstention rates ranging from 0% (CMV and SARS-CoV-2) to 16% (Parvovirus). See Figure 2B and Table 1 for details.

CMV and SARS-CoV-2 classifiers had the strongest performance. CMV dramatically reshapes the T-cell repertoire, and we find 50,217 TCRs in 79 HLA-COclusters specifically associated with CMV. In SARS-CoV-2, samples were acquired before October 2021 and thus were likely all either exposed relatively recently or never exposed/vaccinated, a bygone milieu which necessarily contributes to the very strong performance. The *T. gondii* classifier is remarkably strong given the low prevalence of infection in both the population and our training and holdout donors, suggesting a great impact of *T. gondii* on the T-cell repertoire. The weakness of the Parvovirus classifier at high specificity is likely because Parvovirus is neither a chronic infection, like the herpesviruses (which continually present antigens after initial infection), nor recently acquired in all positive samples, like SARS-CoV-2.

In HSV-2, we separated our training and holdout sets of repertoires by self-reported race, with White donors used for training and Black donors held out for evaluation. This created a meaningful biological difference between the training and holdout sets due to the greater HLA diversity among Black donors. We made this same HSV-2-labeled dataset available for AIRR-ML-25, a public machine learning competition whose results will be published in Nature Methods (35). The strongest competitor’s model achieved AUROC 0.67 with no abstention. That result is not directly comparable with our classifier’s AUROC of 0.95, which abstained on 12 holdout repertoires (12%) due to lack of HLA allele intersection with the relevant ECOcluster. To compare the two more directly, we performed 1,000 assignments of scores randomly chosen from other samples to the 12 abstained samples; median AUROC among those 1,000 predictions on all 100 samples was 0.90. With the important caveats that we had prior access to the labeled repertoires that the competitors lacked and more time to build our models, this result demonstrates the power of the ECOclusters approach to exposure classification.

#### 3.3.3 Model score bimodality in tens of thousands of repertoires

To assess the separation between cases and controls in a broader context, we calculated 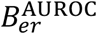 values for each indication on 31,611 repertoires from the previously described T-DETECT cohort (36), which was used in the creation of ECOclusters. For all indications, we observed a bimodal distribution of scores, representing exposed (high) and unexposed (low) modes and ranging from weakly (Parvovirus) to starkly (CMV) bimodal (Figure 2C). We interpret the weak bimodality in Parvovirus to be due to the acute nature of Parvovirus infection, with time since most recent antigen exposure ranging widely among previously exposed donors.

#### 3.3.4 Diagnostic models using multiple ECOclusters

Two of these classifiers, EBV and Parvovirus, make use of multiple ECOclusters, each of which has HLA-COclusters whose breadth is a sensitive and specific predictor of positive label. To investigate why, we assembled a database of TCR-pHLA pairs derived from the public databases VDJDB (30), IEDB (31) and McPAS(32), and intersected the TCRs with ECOclusters.

Three of the four EBV-associated ECOclusters have HLA-COclusters associated with HLA allele A*02:01 and containing TCRs associated with EBV antigens in public databases. In the first, the majority of database-intersecting TCRs are associated with peptide GLCTLVAML from BMLF1 (an early lytic-phase protein involved in mRNA export and processing). In the second, the majority peptide is YVLDHLIVV from BRLF1 (an immediate-early transcription factor that initiates lytic replication), and in the third FLYALALLL from LMP2 (a latent membrane protein expressed during persistent infection). These three EBV-associated ECOclusters may represent immune responses to distinct stages of the viral life cycle that occur at different times in different people.

#### 3.3.5 ECOclusters enriched for TCRs responding to peptides from the same taxon

Following methods used in gene set enrichment analysis, we used a hypergeometric test to identify ECOclusters statistically enriched for TCRs known to respond to peptides from the same taxon (see Methods). We used this approach separately with our database of TCR-pHLA pairs derived from public databases and a database derived from experiments using our MIRA assay.

Two ECOclusters used by the EBV classifier were also associated with EBV by intersection with MIRA and a third via intersection with public databases. Two ECOclusters used by the Parvovirus ECOcluster also gained Parvovirus associations from MIRA. One ECOcluster was associated with Influenza using both databases and additionally with SARS-CoV-2 using the public database. Public databases associated two more ECOclusters with *M. tuberculosis*, while MIRA associated additional ECOclusters with Adenovirus D, Influenza, Coxsackievirus, Polio virus (potentially driven by vaccination) and HHV-6B.

The ECOcluster used by the CMV classifier (“the CMV ECOcluster”) also has large intersection with CMV MIRA TCRs (656) and public database CMV TCRs (239). However, due to the large size of the CMV ECOcluster (50,217 TCRs) and the large amount of MIRA and public CMV data, these large intersections failed to achieve significance via our test.

#### 3.3.6 The CMV ECOcluster in detail

The CMV ECOcluster comprises 50,217 TCRs: 5,323 TCRs from 33 HLA-COclusters associated with Class I HLA alleles and 47,124 TCRs from 58 HLA-COclusters associated with Class II alleles (see Supplementary Data).

Considering *B*_*hr*,_ (HLA-COcluster breadth) calculated on *all* labeled repertoires in the CMV holdout set, without regard to donor HLA, AUROCs for the 91 HLA-COclusters ranged from 0.45 to 0.80 (median: 0.55). 45 of 91 HLA-COclusters had at least three positive-labeled and three negative-labeled repertoires inferred to have the HLA allele with which they are associated. AUROCs on those HLA-restricted repertoires ranged from 0.52 to 1.0 (median: 0.92). All but three HLA-COclusters had dramatically higher AUROC within the HLA-restricted repertoires, illustrating the necessity of accounting for HLA type when comparing TCRs between repertoires (Figure 3A, backing data in Supplementary Data).

**Figure 3.**
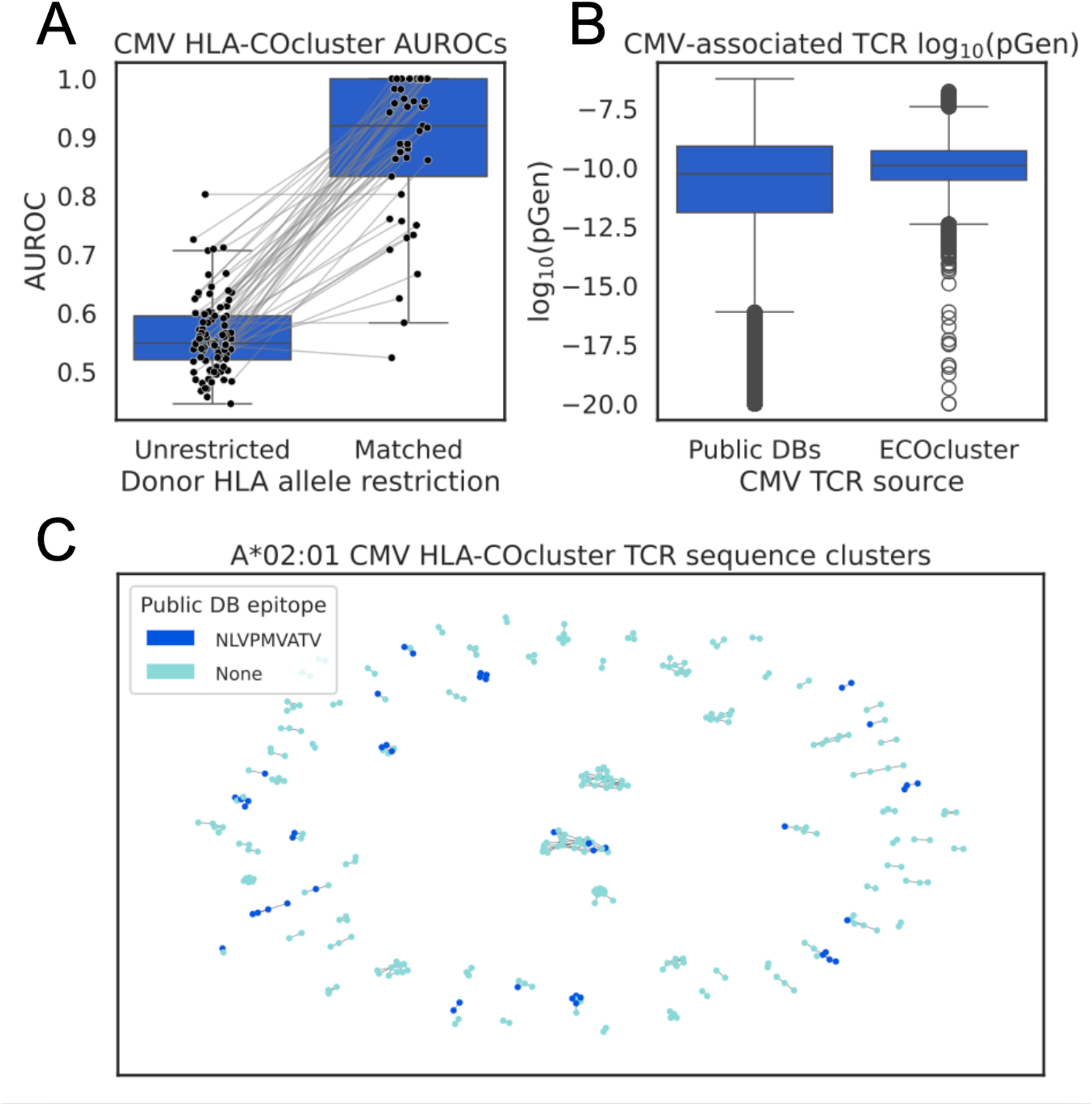
The CMV ECOcluster. A. AUROCs describing *B*_*hr*,_ on HLA-COclusters in the CMV ECOcluster as a classifier of CMV status. All but 3 HLA-COclusters have dramatically higher AUROC when considering only donors inferred to have the associated HLA allele (“Matched”, 45 HLA-COclusters) than when considering all donors (“Unrestricted”, 91). B. TCRs in the CMV ECOcluster have a narrower range of generation probability (pGen) than CMV-associated TCRs in public databases, with higher median pGen. C. 311 of the 540 CMV ECOcluster TCRs associated with A*02:01 have at least one sequence neighbor with a single CDR3 amino acid substitution. Dark blue dots: TCRs associated with CMV peptide NLVPMVATV in public databases. Edges: sequence neighbors.

CMV is well-studied, and our compilation of public TCR databases contains 23,318 TCRs annotated as responding to CMV antigens spanning a pGen range of 2.2e-17 to 1.4e-8 (5th-95th percentile; median: 5.4e-11). The CMV ECOcluster TCRs span a pGen range of 3.3e-12 to 3.7e-9 (median: 1.3e-10). While many public-database CMV TCRs are vanishingly unlikely to be observed again, and therefore of limited utility in annotating new repertoires, the CMV ECOcluster TCRs are public by definition (Figure 3B).

Though HLA-COclusters are defined on TCR co-occurrence, not TCR sequence, we very often observe clusters of sequence-similar TCRs within the same HLA-COcluster. Using a simple definition of a TCR sequence cluster (a connected component of the graph defined by pairs of TCRs with the same V and J gene whose CDR3 sequences differ by a single amino acid substitution), 311 of the 540 TCRs in the CMV HLA-COcluster associated with A*02:01 are members of 79 sequence clusters ranging in cardinality from 2 to 26 TCRs (median: 3). Each sequence cluster notionally represents a TCR binding solution to a single pHLA (Figure 3C).

60 of the 540 TCRs in the CMV HLA-COcluster associated with A*02:01 are present in IEDB, VDJDB and/or McPAS, associated with CMV antigens. 58 are associated with CMV peptide NLVPMVATV and two with VLEETSVML. 46 NLVPMVATV TCRs are members of 21 sequence clusters, 16 of which also contain 50 TCRs not in public databases. Known peptide associations can potentially be propagated within sequence clusters, though confident propagation would require an algorithm proven to cluster TCRs that share pHLA binding.

### 3.4 Typical repertoires contain many ECOcluster TCRs

To investigate the proportion of a typical TCR repertoire annotated by ECOclusters, we assembled 2,000 repertoires from the tdetect_covid, cro_deep_v4b and cro_ultradeep_v4b cohorts that were held out from the construction of our HLA allele-associated TCR database and ECOclusters. Repertoires had between 53,008 and 645,051 unique productive rearrangements (median: 311,524; 5^th^-95^th^ percentile: 121,495.9-545,447.9). We inferred HLA type as described for each repertoire and intersected TCRs from every HLA-COcluster with each repertoire inferred to have its associated HLA allele.

These 2,000 repertoires had between 415 and 16,697 such HLA-matched ECOcluster TCRs (median: 3,327.0; 5^th^-95^th^ percentile: 1,950.8-11,816.1), representing breadths (proportions of repertoire TCRs in any ECOcluster) ranging from 0.001 to 0.041 (median: 0.020; 5^th^-95^th^ percentile: 0.010-0.032) and depths (sums of TCR frequencies) ranging from 0.003 to 0.250 (median: 0.030; 5^th^-95^th^ percentile: 0.014-0.074). Thus, ECOcluster TCRs represent a median of 2.0% of the TCRs in these repertoires and 3.0% of the templates (Figure 4A).

Considering only ECOclusters with a known exposure association (using all methods described above, Figure 4B), repertoires had between 2 and 2,053 total HLA-matched TCRs (median: 455.0; 5^th^-95^th^ percentile: 137.0-1,041.2), representing breadths ranging from 0.000 to 0.007 (median: 0.001; 5^th^-95^th^ percentile: 0.0.001-0.004) and depths ranging from 0.000 to 0.220 (median: 0.004; 5^th^-95^th^ percentile: 0.001-0.037).

**Figure 4.**
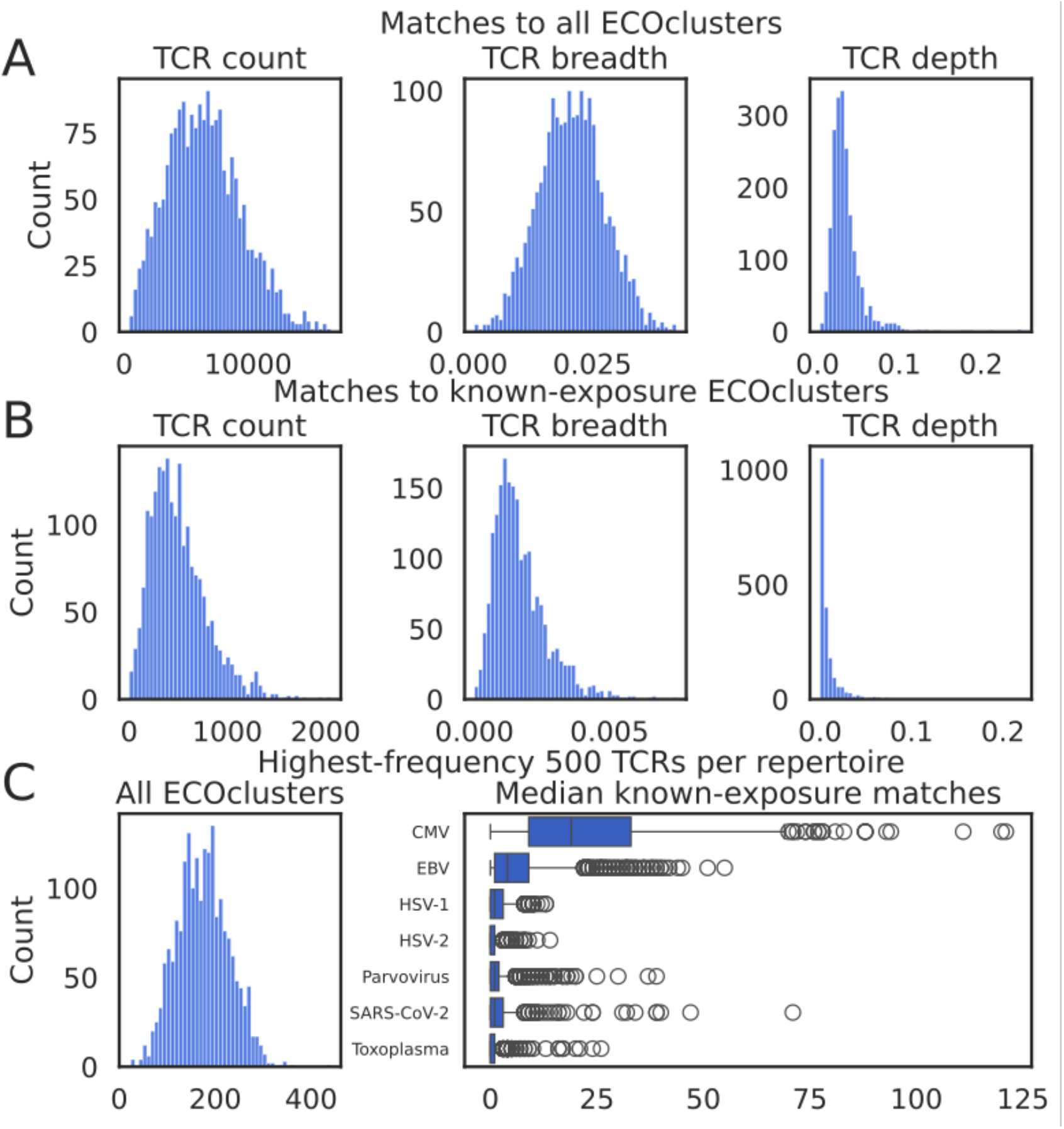
ECOcluster TCR representation in 2,000 repertoires not used to construct ECOclusters. A. Distributions of per-repertoire counts, breadths and depths of HLA-aware matches between repertoire TCRs and ECOclusters. B. As A, but considering only ECOclusters associated with known exposures. C. Representation of ECOcluster TCRs among the highest-frequency 500 TCRs per repertoire. Left: counts of HLA-aware ECOcluster-matches; right: median exposure-associated ECOcluster highest-frequency TCRs per inferred exposure-positive repertoire.

ECOcluster TCRs tend to be well represented among the highest-frequency TCRs in a repertoire (Figure 4C). HLA-matched ECOcluster TCRs represented between 24 and 441 of the 500 highest-frequency TCRs in these repertoires (median: 174.0; 5^th^-95^th^ percentile: 89.0-266.0). The CMV ECOcluster was a major contributor to repertoires inferred to have CMV infection. A median of 19.0 of the highest-frequency TCRs per predicted CMV+ repertoire were members of the CMV-associated ECOcluster, eclipsing the other exposures with diagnostic models (EBV: 4.0; SARS-CoV-2, Parvovirus, HSV-1: 1.0; HSV-2, *T. gondii*: 0.0). While median breadth on the CMV ECOcluster among inferred CMV+ donors was just 0.0009 (5^th^-95^th^ percentile: 0.0003-0.0022), median depth was 0.006 (5^th^-95^th^ percentile: 0.0005-0.055), with the highest depth being 0.219: in one repertoire 21.9% of the TCR templates were CMV ECOcluster members, illustrating the radical expansion of TCR clones responding to CMV.

## 4 Discussion

ECOclusters (TCR clusters derived from co-occurrence in tens of thousands of human immune repertoires) putatively represent the public T-cell responses to many prevalent immune exposures, including viral and bacterial infections, vaccines and medications. ECOclusters represent responses to exposures at least as rare as *T. gondii* (^∼^11% prevalence in the U.S.) and at least as prevalent as EBV (^∼^90%), but the exposures driving most ECOclusters remain unknown.

Our co-clustering approach has advantages over other means of annotating TCR specificity. TCRs (TCR β chains) experimentally determined to bind a pHLA could be part of a TCR α-β pair that binds a different pHLA presented by another prevalent immune exposure. By contrast, TCRs in an exposure-associated ECOcluster are unlikely to bind antigens presented during other prevalent exposures: if they did, they would not cluster by occurrence with other exposure-specific TCRs. TCR annotations derived from ECOclusters can therefore be made with greater confidence than annotations derived only from TCR-pHLA binding experiments.

An earlier version of the CMV-associated ECOcluster than the one described in this work, made publicly available in our earlier preprint, has already proven useful to researchers. Grabauskas et al. first validated that CMV ECOcluster’s sensitivity and specificity as a biomarker of the CMV T-cell response. Then, by intersecting the CMV ECOcluster TCRs with single-cell T cell RNA data, they demonstrated that the well-known expansion of CD8^+^ and CD4^+^ TEMRA cells with CMV infection is reflected in T cells with CMV-specific TCRs. They also reported the novel finding that CMV-specific clones are enriched for GZMK^+^ CD8^+^ and CD4^+^ Th1 T cells (21).

Though ECOclusters necessarily represent responses to prevalent exposures, multiple researchers have applied them to studies of autoimmune disorders, using an earlier version of ECOclusters encompassing fewer HLA alleles and built from fewer TCR repertoires. El Abd et al. demonstrated that donors with Primary Sclerosing Cholangitis (PSC) have more EBV-responding ECOcluster TCRs and that individual TCRs elevated in PSC are part of the public response to EBV (22). We previously used CMV- and EBV-associated ECOclusters to show that CMV and EBV TCRs were significantly elevated in Celiac+ donors vs. controls, but only among donors who had had the respective viral exposures (23). An earlier version of the CMV ECOcluster was used to establish a strong relationship between CMV and a rare neuroimmunological disease (publication forthcoming).

The knowledge that a TCR is associated with a prevalent immune exposure can shed light on that TCR’s role in an immune repertoire. We observe many ECOcluster-member TCRs (including CMV) in publicly available tumor single-cell RNA experiments (uTILity, https://github.com/ncborcherding/utility). While possibly binding a tumor-related antigen, those TCRs are more parsimoniously understood as bystanders. The fraction of tumor-infiltrating lymphocytes (TILs) that are tumor-responding can be an important prognostic indicator (37), and so understanding the composition of TILs can have implications for clinical outcomes.

The T-cell repertoire is a ledger of prior exposures that has remained largely uninterpretable. ECOclusters represent a step toward deciphering that ledger, with applications across many areas of medicine.

## 6 Author Contributions

DHM and SW contributed concept and algorithm development, computational modeling and writing. JG provided concept development and oversight. AJM and RT contributed experimental work and writing. BG and RT contributed laboratory assays and oversight. TH, ZY, JRG and RB contributed laboratory assays. MTN and RE contributed concept development. KD, EJO and JO contributed data analysis. HCH, AS, BH, MK and HSR contributed oversight.

## 7 Acknowledgments

We thank Monera Alrukhayes and colleagues at Kuwait University, who collected samples from which sequencing data was used for this study. We thank H. Jabran Zahid for initial development of models predicting HLA allele status from TCR repertoires and for contributions to the ideas embodied in this work.

## 8 Ethics Statement

The studies involving humans were approved for each cohort as cited in the earlier studies referenced using the same cohorts, except the control_bloodworks cohort (not used in previous work), collected under Bloodworks Research Donor Collection Protocol BT001.

## 9 Data Availability

The TCRs in the CMV ECOcluster, with their HLA-COcluster and HLA associations, are provided as a supplementary zip-compressed tab-separated-value file whose columns are described in Supplementary Materials.

