## Supplementary Materials for "Immune signatures of common exposures through co-occurrence of T-cell receptors in tens of thousands of donors"

### *Supplementary Material*

#### 1 Supplementary Methods

##### 1.1 Constructing classifiers for HLA allele status

We built classification models for each of 220 HLA alleles (where an “HLA allele” can be an individual allele, a class II DP/DQ heterodimer or a p group – a set of HLA alleles with identical antigen binding domain protein sequences) with a modified version of the methodology previously described. For each HLA allele *a*:

1. Split the “case”- and “control”-labeled repertoires with respect to *a* randomly 80:20 into training and holdout. We use a fixed split across all HLA alleles.
2. Identify a set of candidate TCRs overrepresented in cases by Fisher’s Exact Test, using a *p*-value threshold hyperparameter determined in 5-fold cross-validation.
3. Use the “L1LR” method described previously to retain only the candidate TCRs that associate most strongly with the HLA being modeled
4. Train a logistic regression classifier for allele presence with two features:

   $F_l = \left[\log\left(1 + \sum_j w_j\, c_{rj}\right), \log(1 + N_r)\right]$, where $c_{rj}$ is the *convergent recombination count* (the number of unique nucleotide rearrangements for associated TCR *j*) and $N_r$ is the total unique rearrangements in repertoire *r*.

5. Set decision thresholds for calling high-confidence positives (at 95% precision) and negatives (at 95% recall on positive labels).
6. Retain models with ≥ 95% precision and ≥ 10 positive and negative calls in cross validation and/or holdout.

##### 1.2 Associating public TCRs with HLA alleles

We used an empirical method to calculate FDR. We first performed random label assignments on a grid of parameters to estimate the number of significant TCRs at a given FET *p*-value threshold expected at random, using our tdetect_covid and cmv_cohort_1 cohorts. We performed 100 label permutations for each combination of parameter values, varying the number of cases/controls from 20 to 6000 (to account for the large variation in the prevalence of HLA alleles). We observed that this quantity, as a function of the number of cases and controls, follows a power distribution. We fit a single linear model to estimate the expected number of significant TCRs at a given *p*-value threshold from the logarithm of the number of cases, the logarithm of the number of controls, and the average total unique rearrangements in the repertoires under consideration:

$$f = \frac{\text{avg\_depth}}{\text{ad}_0}$$

$$m = \alpha_1 \log_{10}(n_{\text{cases}}) + \alpha_2 \log_{10}(n_{\text{controls}}) + \alpha_3 f + \alpha_0,$$
$$b = \beta_1 \log_{10}(n_{\text{cases}}) + \beta_2 \log_{10}(n_{\text{controls}}) + \beta_3 f + \beta_0.$$

$$\log_{10} \hat{C}_i = m \log_{10}(q_i) + b$$

Where avg_depth is the average number of productive rearrangements in the samples under consideration and $ad_0$ is the average number of productive rearrangements of the samples used to fit the model. $q_i$ is the $i$-th FET $p$-value (for which we wish to derive an FDR).

We then derive a log FDR value:

$$C_i = \sum_{j=1}^{N} \mathbf{1}\{q_j \leq q_i\}$$

$$\log_{10}(\text{FDR}_i) = \min\{0, \log_{10} \hat{C}_i - \log_{10}(C_i)\}$$

Here, $C_i$ is the observed number of FET $p$-values less than or equal to $q_i$, and $\hat{C}_i$ is the model-predicted number at that threshold. The FDR is the ratio of these two quantities.

##### 1.3 Serological labeling

For EBV, Parvovirus, HSV-1, HSV-2 and *T. gondii*, we derived new serological labels on previously acquired samples. A multiplexed serological testing method was developed in house using U-PLEX Development Pack from Meso Scale Discovery (MSD). Purified antigens (recombinant VCA p18 and EBNA-1 proteins for EBV, recombinant HSV-1 gG protein, recombinant HSV-2 gG protein and *T. gondii* antigen were purchased from Meridian Life Science. Parvovirus B19 VLP/VP1/VP2 Co-Capsid Recombinant protein was purchased from Raybiotech) were biotinylated at optimized biotin-to-protein ratios that generated biotinylated proteins with 1-3 biotin(s) per molecule.

Biotinylated antigens were coated on the plate simultaneously at optimized concentrations onto different spots via linker provided by MSD. After washing off the unbound antigens, sera samples diluted to optimized concentration with assay diluent were applied to the plate. Antibodies in the serum that recognize the plate bound antigens were detected by a sulfo-tag labeled anti-human IgG antibody. The signal level of each spot is in direct correlation with the amount of antigen-specific antibodies in the serum sample. A positive control that contains antibodies against all the antigens in the panel, a negative control that does not have detectable antibodies against any of the antigens in the panel and two cutoff samples that contain threshold level of antibodies against the antigens in the panel were run on each plate. The multiplexed serological testing method was validated using clinically labeled serum samples and using commercially available ELISA kits.

We normalized MSD signal in two steps to remove variation in background signal among (1) wells and samples, and (2) MSD plates. First, within each well of an MSD plate, we used the mean signal of spots without antigens as a measure of background signal and subtracted it from the signal of spots with bound disease antigens. We ran each sample in three wells and took the mean of the three background-adjusted signal values for each disease. Second, to remove variation among MSD plates, we included a cutoff sample in three wells of every plate and calculated the normalized signal, *S*, for each disease for each sample; for each antigen, we divided the mean background-adjusted signal of every sample on the plate by the mean background-adjusted signal of the cutoff sample. The signal of the cutoff sample was always greater than the background signal, so the denominator of *S* was always positive, but for some samples with low signal for a disease, the numerator (and *S*) was negative.

##### 1.4 Statistical methods for identifying high-confidence serological labels

We observed a bimodal distribution of log *S* for each disease, approximating a mixture of two Guassians. Accordingly, we modeled log *S* as a mixture of two univariate Gaussian distributions, assuming the component distributions with lower and higher signal represented controls and cases, respectively. When fitting the mixture model, we ignored all samples with negative *S;* this ranged from 1-6% of the samples among the diseases. After fitting the means, variances, and mixture proportion of the mixture model using all samples with positive *S*, we used Bayes' rule to calculate the probability each sample was a case given its value of log *S*. When calculating this probability for samples with negative *S*, we used the smallest positive *S* among the samples for the disease. For model training and evaluation, we considered samples with probability less than 0.01 and greater than 0.99 as high-confidence controls and cases, respectively.

For the EBV labels, as described above, we had labels and confidence estimates for two antigens, VCA and EBNA1. We used VCA as our primary indicator of donor EBV status. However, a small number of donors had a confident negative label for VCA but also had a label for EBNA1 that was insufficiently confidently negative. To ensure EBV label quality, with no clear approach to combining two disagreeing labels into one, we omitted samples with confident negative labels for VCA and >0.1 posterior probability of positive label for EBNA1.

#### 2 Supplementary Results

##### 2.1 Models predicting donor status with respect to 220 HLA alleles from TCR repertoires

We previously published methods for imputing HLA type from the TCRβ repertoire, resulting in 135 high-precision HLA models (24). Here, we extend this approach to 85 more HLA alleles, trained and evaluated on TCRβ repertoires from 1,810 more HLA-genotyped individuals sourced from more ethnically and racially diverse populations (Supplementary Figure 1A).

We model presence of 220 HLA alleles at a minimum of 95% precision; 11 models represent two or more alleles modeled collectively as a P group (distinct HLA alleles with identical antigen binding domain protein sequences) and 82 represent Class II heterodimers rather than individual alleles (Supplementary Table 2). We were able to construct high-precision models for 70% of the HLA alleles observed at 1% or greater prevalence in our training data (Supplementary Figure 1B), with some models constructed using as few as 30 training positive labels. Overall, our models have a median precision of 97%, median recall of 74% and median negative predictive value of 99.9%.

##### 2.2 Eight million public TCRs associated with HLA alleles

We used our 220 HLA models to computationally HLA type a much larger cohort of 34,793 individuals, consisting of subjects used to train the HLA models described above and 28,853 additional subjects with unknown HLA type (Supplementary Table 1). We held out from this process repertoires also held out from HLA model training and repertoires later used to evaluate diagnostic model performance.

To construct a large database of HLA allele-associated TCRs, we performed Fisher's Exact Test (FET) for each public TCR against each imputed HLA allele label to statistically associate HLA alleles with TCRs. We then used a lowest False Discovery Rate (FDR) approach to uniquely associate each TCR with a single HLA allele, yielding a database of 8,618,285 HLA allele-associated TCRs. 92% of TCRs were associated with Class II HLA alleles and 8% with Class I alleles.

To be associated with an HLA allele, a TCR's pGen must necessarily be high enough to be observable in multiple donors, but not so high as to be observed in many donors regardless of their response to their cognate antigen from a prevalent exposure. Accordingly, the pGen values of the HLA allele-associated TCRs spanned 3.6e-12 to 5.6e-9 (5th-95th percentile), with median 1.7e-10, while the same number of TCRs randomly sampled from the same repertoires spanned the much wider range 7.7e-15 to 1.4e-8, with median 9.5e-11 (Supplementary Figure 2).

#### 3 Supplementary Figures and Tables

##### 3.1 Supplementary Figures

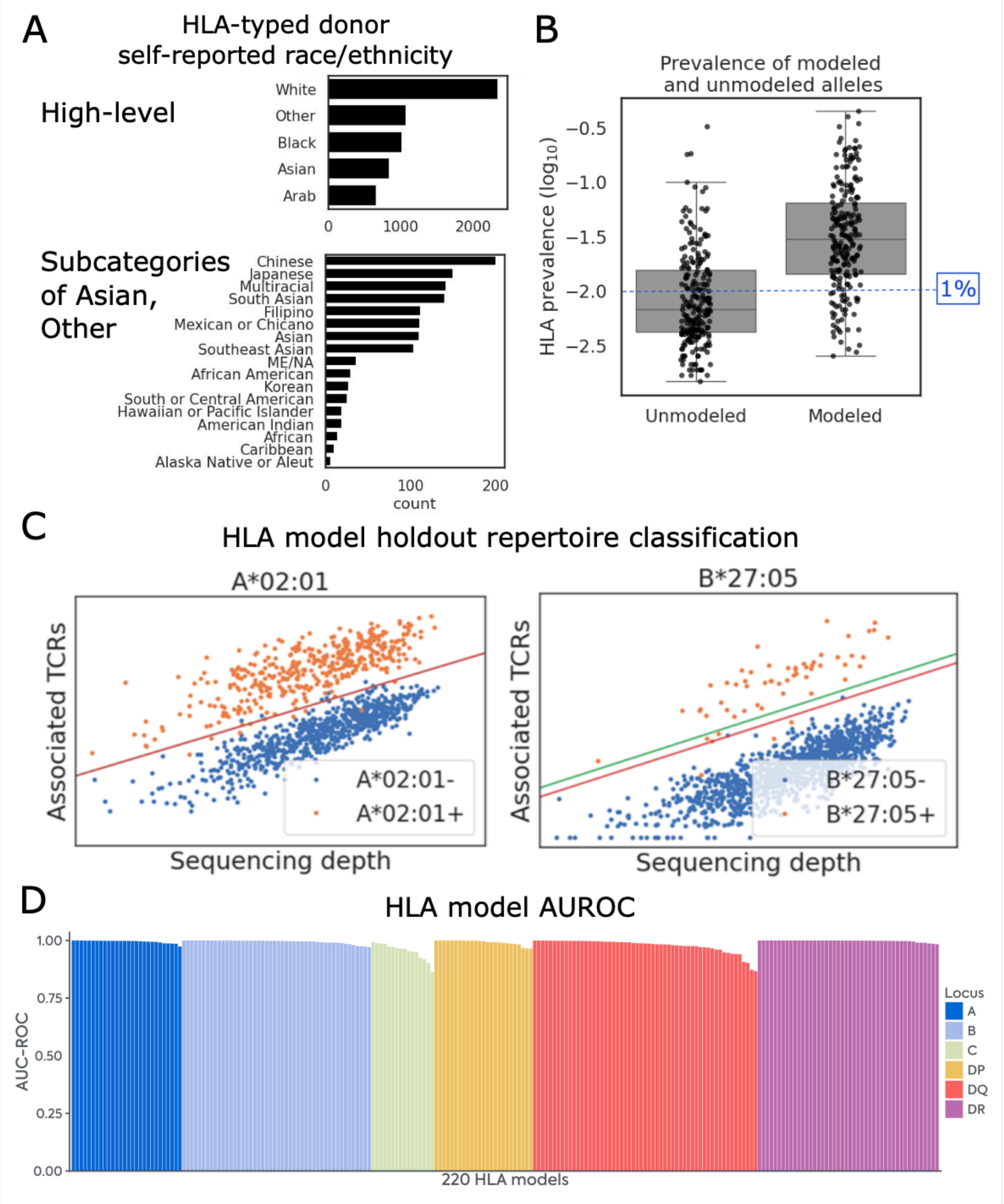

**Supplementary Figure 1.** Classifier models for 220 HLA alleles. A. Self-reported race/ethnicity of donors used to train and evaluate HLA models. Top bar chart shows gross split of donor demographics and lower bar chart shows subdivision of Asian/Other categories. B. Prevalence of all HLA alleles observed in training/evaluation data, shown in log scale, split by whether alleles are modeled. We were able to construct high-precision models for 70% of the HLA alleles observed at 1% or greater prevalence in our training data, including alleles with as few as 30 training cases C. Models classify repertoires as HLA allele+ or allele- using convergent recombination count of allele-associated TCRs (vertical axis) and total productive rearrangements (sequencing depth, horizontal axis). Positive (green) and negative (red) thresholds define positive, negative and indeterminate calls. For A*02:01, these thresholds converge (no indeterminate calls). D. AUROCs of the 220 models across each locus, sorted by descending AUROC.

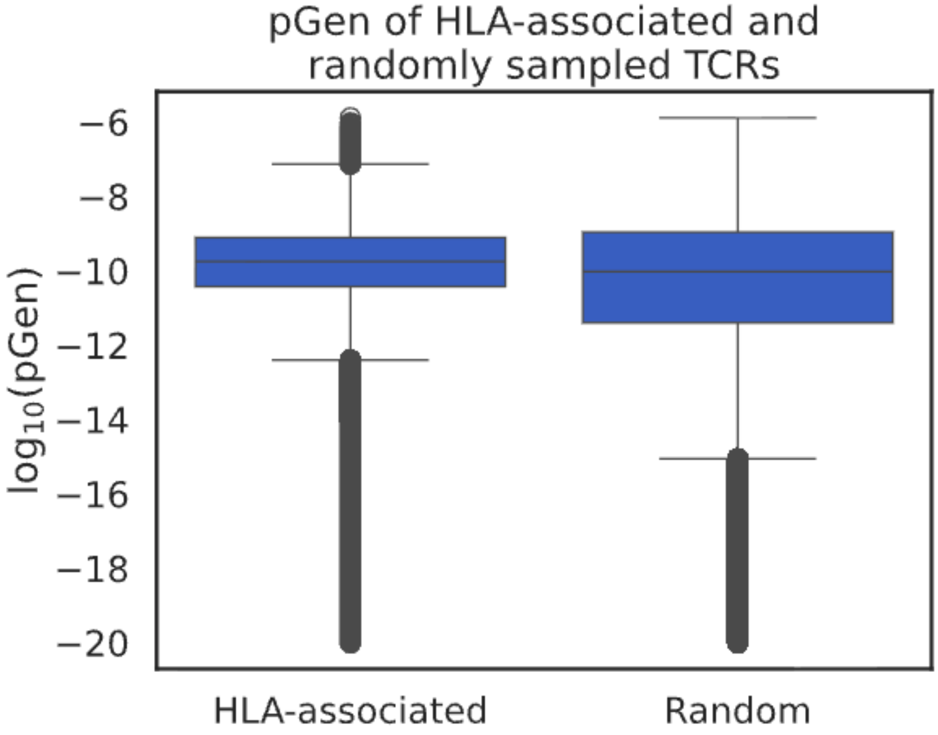

**Supplementary Figure 2.** TCR V(D)J generation probability (pGen, as computed by OLGA). pGen of 8,618,285 HLA allele-associated TCRs spanned 3.6e-12 to 5.6e-9 (5th-95th percentile), with median 1.7e-10, while the same number of TCRs randomly sampled from the same repertoires spanned the much wider range 7.7e-15 to 1.4e-8, with median 9.5e-11.

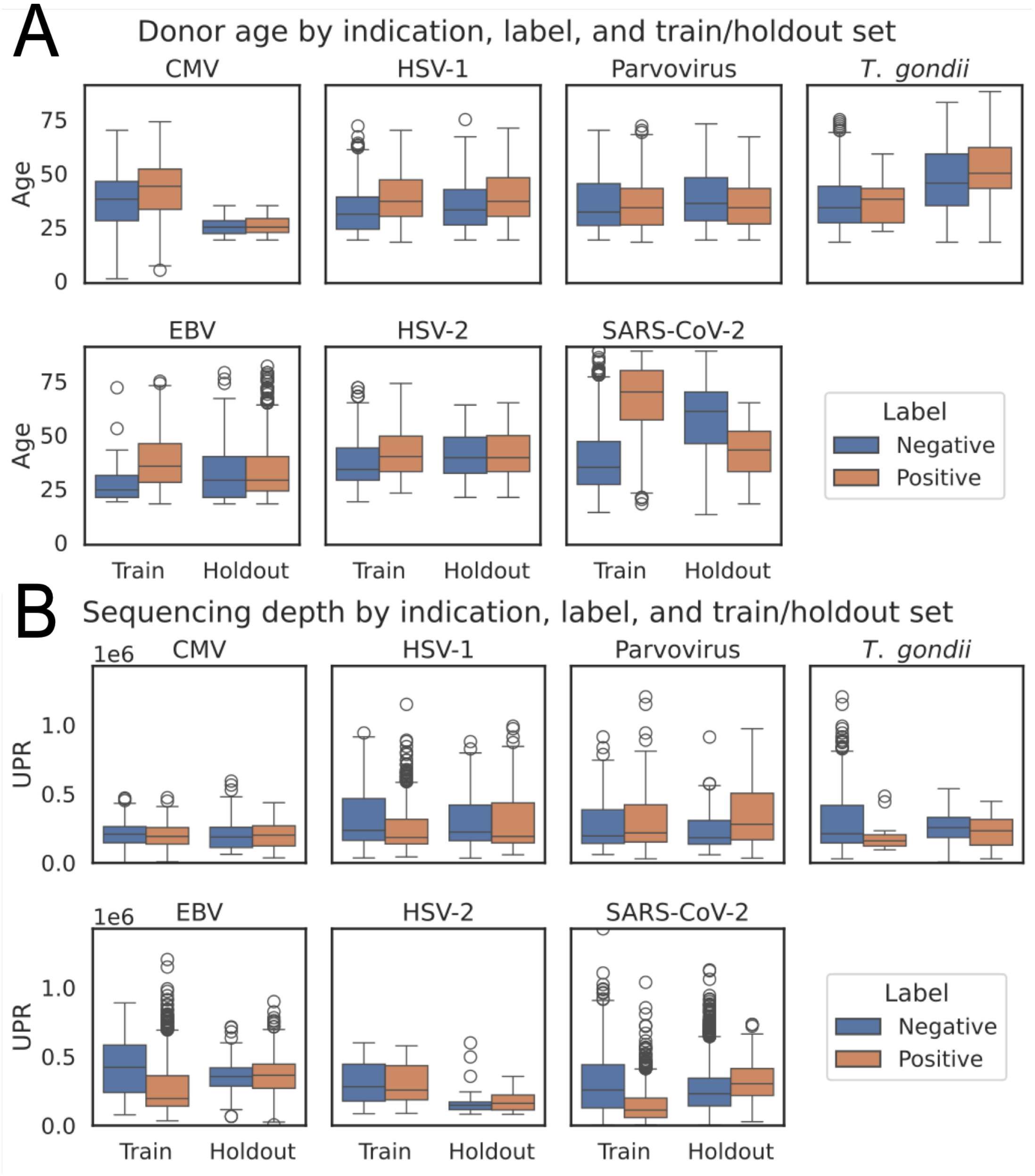

**Supplementary Figure 3.** Distributions of positively and negatively labeled samples for each indication, within train and holdout sets. A. Donor age. B. Unique productive rearrangements (UPR).

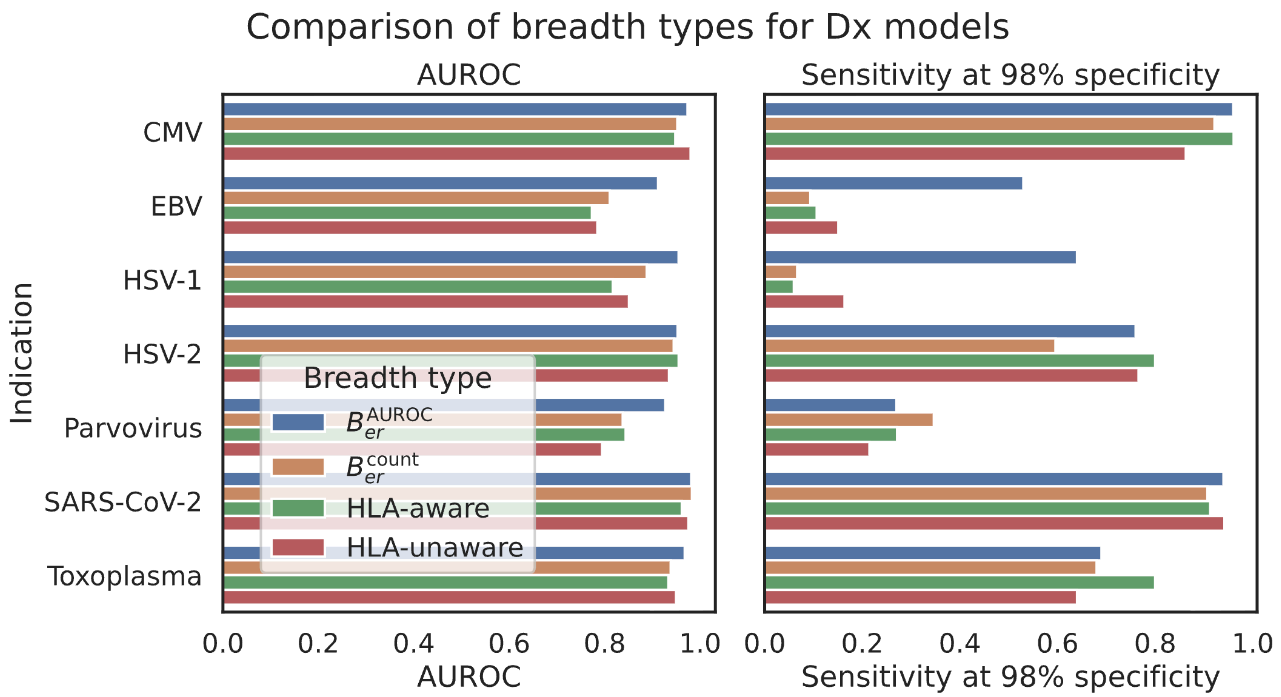

**Supplementary Figure 4.** Statistics describing classifier performance for each indication using four different breadth measures on the same ECOclusters: $B_{er}^{\text{AUROC}}$ and $B_{er}^{\text{count}}$, as described in Methods, HLA-aware breadth (matching TCRs in repertoires only if the repertoire's donor is inferred to have their associated HLA allele) and HLA-unaware breadth (matching TCRs regardless of donor HLA). The more complex weighting of $B_{er}^{\text{AUROC}}$, learned from training samples, was particularly helpful in classifying two of the "hardest" indications, EBV and HSV-1. Left: AUROC. Right: sensitivity at 98% specificity.

#### 3.2 Supplementary Tables

**Supplementary Table 1.** Descriptions of sample cohorts. Breakdown by self-reported sex and age, 5th-95th percentile ranges of TCR rearrangement counts and age (when known), and whether the cohorts were used in building HLA allele modeling, ECOcluster construction and exposure classifier training and holdout sets. From all cohorts, individual samples used in HLA model and exposure classifier holdout sets were excluded from HLA modeling and ECOcluster construction.

| Cohort | Repertoires | Male | Female | Unknown sex | UPR | Age | HLA models | ECOclusters | Classifier training | Classifier holdout |
|---|---|---|---|---|---|---|---|---|---|---|
| cmv_cohort_1 | 666 | 345 | 297 | 24 | 64582.0 - 331711.5 | 13.7 - 61.0 | Yes | No | Yes | No |
| cmv_cohort_2 | 120 | 47 | 73 | 0 | 79591.2 - 424576.8 | 19.0 - 34.0 | Yes | No | No | Yes |
| control_bloodworks | 5020 | 2218 | 2610 | 192 | 140708.1 - 658344.5 | unknown | Yes | Yes | No | Yes |
| covid_dls | 628 | 326 | 300 | 2 | 25985.9 - 375771.4 | 33.2 - 89.0 | Yes | No | Yes | No |
| covid_isb | 444 | 203 | 239 | 2 | 42363.8 - 530032.3 | 26.0 - 83.8 | Yes | No | Yes | No |
| covid_koelle | 682 | 174 | 213 | 295 | 538.5 - 479990.2 | 24.3 - 70.0 | Yes | No | No | No |
| covid_kuwait | 621 | 0 | 0 | 621 | 24471.0 - 545588.0 | unknown | Yes | No | No | No |
| cro_deep_v4 | 1871 | 761 | 1110 | 0 | 50605.5 - 166599.5 | 21.0 - 62.0 | Yes | Yes | No | No |
| cro_deep_v4b | 1103 | 669 | 434 | 0 | 84456.9 - 280986.9 | 21.0 - 60.9 | Yes | Yes | Yes | Yes |
| cro_ultradeep_v4 | 120 | 60 | 60 | 0 | 222281.4 - 586300.0 | 20.0 - 66.0 | Yes | Yes | No | No |
| cro_ultradeep_v4b | 2083 | 782 | 1301 | 0 | 143553.8 - 702539.4 | 21.0 - 67.0 | Yes | Yes | Yes | Yes |
| lyme_jh | 446 | 244 | 202 | 0 | 48297.5 - 150451.0 | 25.0 - 73.0 | Yes | No | No | No |
| tdetect_covid | 32276 | 15237 | 16899 | 53 | 143089.3 - 543953.8 | 26.0 - 71.0 | No | Yes | No | No |

**Supplementary Table 2.** List of 220 modeled HLA alleles. "+" between two alleles indicates a Class II heterodimer. Cases and controls across all training and holdout data are given. For some P group models, it was also possible to build high-precision models for individual alleles in the P group, but we found those models were essentially indistinguishable (listed in the 'Alleles combined' column).

| HLA model | Cases | Controls | Alleles combined | HLA model | Cases | Controls | Alleles combined |
|---|---|---|---|---|---|---|---|
| A*01:01 | 1133 | 4821 | | B*40:02 | 181 | 5773 | |
| A*02:01 | 2053 | 3901 | | B*40:06 | 130 | 5824 | |
| A*02:02 | 109 | 5845 | | B*41:01 | 85 | 5869 | |
| A*02:03 | 69 | 5885 | | B*42:01 | 134 | 5820 | |
| A*02:05 | 136 | 5818 | | B*44:02 | 535 | 5419 | |
| A*02:06 | 148 | 5806 | | B*44:03 | 545 | 5409 | |
| A*02:07 | 124 | 5830 | | B*44:05 | 14 | 5940 | |
| A*02:11 | 31 | 5923 | | B*45:01 | 154 | 5800 | |
| A*03:01 | 1028 | 4926 | | B*46:01 | 162 | 5792 | |
| A*11:01 | 819 | 5135 | | B*47:01 | 18 | 5936 | |
| A*23:01P | 384 | 5570 | A*23:01, A*23:17 | B*48:01 | 68 | 5886 | |
| A*24:02 | 1096 | 4858 | | B*49:01 | 187 | 5767 | |
| A*24:07 | 109 | 5845 | | B*50:01 | 196 | 5758 | |
| A*25:01 | 130 | 5824 | | B*51:01 | 586 | 5368 | |
| A*26:01 | 303 | 5651 | | B*52:01 | 229 | 5725 | |
| A*29:02 | 286 | 5668 | | B*53:01 | 284 | 5670 | |
| A*30:01 | 288 | 5666 | | B*55:01 | 160 | 5794 | |
| A*30:02 | 207 | 5747 | | B*57:01 | 242 | 5712 | |
| A*31:01 | 358 | 5596 | | B*57:03 | 81 | 5873 | |
| A*32:01 | 308 | 5646 | | B*58:01 | 259 | 5695 | |
| A*33:01 | 126 | 5828 | | B*78:01 | 21 | 5933 | |
| A*33:03 | 359 | 5595 | | C*01:02 | 608 | 5346 | |
| A*34:01 | 66 | 5888 | | C*02:02P | 476 | 5478 | |
| A*34:02 | 81 | 5873 | | C*03:04 | 791 | 5163 | |
| A*36:01 | 72 | 5882 | | C*04:01 | 1412 | 4542 | |
| A*68:01 | 355 | 5599 | | C*05:01 | 573 | 5381 | |
| A*68:02 | 232 | 5722 | | C*06:02 | 914 | 5040 | |
| A*74:01 | 148 | 5806 | | C*07:01P | 1237 | 4717 | |
| B*07:02 | 942 | 5012 | | C*07:02 | 1395 | 4559 | |
| B*07:05P | 159 | 5795 | | C*08:01 | 228 | 5726 | |
| B*08:01 | 807 | 5147 | | C*08:02 | 360 | 5594 | |
| B*13:01 | 72 | 5882 | | C*12:02 | 226 | 5728 | |
| B*13:02 | 188 | 5766 | | C*12:03 | 423 | 5531 | |
| B*14:02 | 259 | 5695 | | C*14:02 | 202 | 5752 | |
| B*15:01 | 448 | 5506 | | C*16:01 | 447 | 5507 | |
| B*15:02 | 106 | 5848 | | C*17:01P | 292 | 5662 | |
| B*15:03 | 147 | 5807 | | C*18:01P | 70 | 5884 | |

| | | | | | | |
|---|---|---|---|---|---|---|
| B*15:10 | 101 | 5853 | | DPA1*01:03+DPB1*02:01 | 1528 | 4426 |
| B*15:16 | 49 | 5905 | | DPA1*01:03+DPB1*02:02 | 82 | 5872 |
| B*15:17 | 62 | 5892 | | DPA1*01:03+DPB1*03:01 | 662 | 5292 |
| B*15:18 | 40 | 5914 | | DPA1*01:03+DPB1*04:01 | 2662 | 3292 |
| B*18:01 | 394 | 5560 | | DPA1*01:03+DPB1*04:02 | 982 | 4972 |
| B*27:04 | 29 | 5925 | | DPA1*01:03+DPB1*06:01 | 116 | 5838 |
| B*27:05 | 249 | 5705 | | DPA1*01:03+DPB1*104:01 | 249 | 5705 |
| B*35:01 | 604 | 5350 | | DPA1*01:03+DPB1*13:01 | 174 | 5780 |
| B*35:03 | 180 | 5774 | | DPA1*01:03+DPB1*15:01 | 53 | 5901 |
| B*35:05 | 93 | 5861 | | DPA1*01:03+DPB1*18:01 | 158 | 5796 |
| B*35:08 | 53 | 5901 | | DPA1*01:03+DPB1*85:01 | 15 | 5939 |
| B*37:01 | 109 | 5845 | | DPA1*02:01+DPB1*01:01 | 671 | 5283 |
| B*38:01 | 149 | 5805 | | DPA1*02:01+DPB1*09:01 | 116 | 5838 |
| B*38:02 | 91 | 5863 | | DPA1*02:01+DPB1*131:01 | 53 | 5901 |
| B*39:01 | 120 | 5834 | | DPA1*02:01+DPB1*13:01 | 265 | 5689 |
| B*39:05 | 33 | 5921 | | DPA1*02:01+DPB1*14:01 | 223 | 5731 |
| B*39:06 | 55 | 5899 | | DPA1*02:01+DPB1*17:01 | 238 | 5716 |
| B*40:01 | 502 | 5452 | | DPA1*02:02+DPB1*01:01 | 511 | 5443 |
| DPA1*02:02+DPB1*05:01 | 545 | 5409 | | DQA1*05:03+DQB1*03:01 | 64 | 5890 |
| DPA1*02:02+DPB1*105:01 | 37 | 5917 | | DQA1*05:05+DQB1*02:01 | 114 | 5840 |
| DPA1*02:02+DPB1*135:01 | 53 | 5901 | | DQA1*05:05+DQB1*02:02 | 118 | 5836 |
| DPA1*02:06+DPB1*05:01 | 43 | 5911 | | DQA1*05:05+DQB1*03:01 | 1037 | 4917 |
| DPA1*02:07+DPB1*19:01 | 45 | 5909 | | DQA1*05:05+DQB1*03:03 | 65 | 5889 |
| DPA1*02:12+DPB1*85:01 | 30 | 5924 | | DQA1*05:05+DQB1*03:19 | 146 | 5808 |
| DPA1*03:01+DPB1*105:01 | 199 | 5755 | | DQA1*05:05+DQB1*05:01 | 146 | 5808 |
| DQA1*01:01+DQB1*02:01 | 96 | 5858 | | DQA1*05:09+DQB1*03:01 | 31 | 5923 |
| DQA1*01:01+DQB1*02:02 | 115 | 5839 | | DQA1*06:01+DQB1*03:01 | 271 | 5683 |
| DQA1*01:01+DQB1*05:01 | 966 | 4988 | | DRB1*01:01 | 645 | 5309 |
| DQA1*01:01+DQB1*05:03 | 42 | 5912 | | DRB1*01:02 | 234 | 5720 |
| DQA1*01:01+DQB1*06:02 | 111 | 5843 | | DRB1*01:03 | 74 | 5880 |
| DQA1*01:01+DQB1*06:03 | 42 | 5912 | | DRB1*03:01 | 1052 | 4902 |
| DQA1*01:02+DQB1*03:01 | 370 | 5584 | | DRB1*03:02 | 159 | 5795 |
| DQA1*01:02+DQB1*05:02 | 435 | 5519 | | DRB1*04:01 | 569 | 5385 |
| DQA1*01:02+DQB1*06:01 | 115 | 5839 | | DRB1*04:02 | 86 | 5868 |
| DQA1*01:02+DQB1*06:02 | 1212 | 4742 | | DRB1*04:03 | 170 | 5784 |
| DQA1*01:02+DQB1*06:04 | 312 | 5642 | | DRB1*04:04 | 281 | 5673 |
| DQA1*01:02+DQB1*06:09 | 204 | 5750 | | DRB1*04:05 | 266 | 5688 |
| DQA1*01:03+DQB1*02:01 | 80 | 5874 | | DRB1*04:07 | 109 | 5845 |
| DQA1*01:03+DQB1*05:01 | 92 | 5862 | | DRB1*04:08 | 37 | 5917 |
| DQA1*01:03+DQB1*05:03 | 46 | 5908 | | DRB1*04:11 | 20 | 5934 |
| DQA1*01:03+DQB1*06:01 | 280 | 5674 | | DRB1*07:01 | 1244 | 4710 |
| DQA1*01:03+DQB1*06:02 | 89 | 5865 | | DRB1*08:01 | 154 | 5800 |
| DQA1*01:03+DQB1*06:03 | 476 | 5478 | | DRB1*08:02 | 105 | 5849 |

| | | | | | | | |
|---|---|---|---|---|---|---|---|
| DQA1*01:04+DQB1*05:01 | 55 | 5899 | | DRB1*08:03 | 121 | 5833 | |
| DQA1*01:04+DQB1*05:03 | 278 | 5676 | | DRB1*08:04 | 153 | 5801 | |
| DQA1*01:04+DQB1*06:02 | 35 | 5919 | | DRB1*08:06 | 18 | 5936 | |
| DQA1*01:05+DQB1*05:01 | 300 | 5654 | | DRB1*09:01 | 357 | 5597 | |
| DQA1*02:01+DQB1*02:01 | 122 | 5832 | | DRB1*10:01 | 210 | 5744 | |
| DQA1*02:01+DQB1*02:02 | 1004 | 4950 | | DRB1*11:01 | 626 | 5328 | |
| DQA1*02:01+DQB1*03:03 | 277 | 5677 | | DRB1*11:02 | 107 | 5847 | |
| DQA1*02:01+DQB1*03:19 | 25 | 5929 | | DRB1*11:03 | 39 | 5915 | |
| DQA1*02:01+DQB1*04:02 | 53 | 5901 | | DRB1*11:04 | 255 | 5699 | |
| DQA1*02:01+DQB1*05:01 | 170 | 5784 | | DRB1*12:01 | 250 | 5704 | |
| DQA1*02:01+DQB1*06:02 | 156 | 5798 | | DRB1*12:02 | 250 | 5704 | |
| DQA1*02:01+DQB1*06:03 | 66 | 5888 | | DRB1*13:01 | 558 | 5396 | |
| DQA1*02:01+DQB1*06:04 | 41 | 5913 | | DRB1*13:02 | 565 | 5389 | |
| DQA1*03:01+DQB1*02:01 | 90 | 5864 | | DRB1*13:03 | 148 | 5806 | |
| DQA1*03:01+DQB1*02:02 | 85 | 5869 | | DRB1*13:04 | 23 | 5931 | |
| DQA1*03:01+DQB1*03:02 | 807 | 5147 | | DRB1*14:01P | 291 | 5663 | DRB1*14:01, DRB1*14:54 |
| DQA1*03:01+DQB1*05:01 | 98 | 5856 | | DRB1*14:02 | 27 | 5927 | |
| DQA1*03:02+DQB1*03:03 | 286 | 5668 | | DRB1*14:04 | 47 | 5907 | |
| DQA1*03:03+DQB1*02:01 | 84 | 5870 | | DRB1*14:06 | 27 | 5927 | |
| DQA1*03:03+DQB1*02:02 | 182 | 5772 | | DRB1*15:01 | 967 | 4987 | |
| DQA1*03:03+DQB1*03:01 | 430 | 5524 | | DRB1*15:02 | 345 | 5609 | |
| DQA1*04:01+DQB1*03:19 | 84 | 5870 | | DRB1*15:03 | 304 | 5650 | |
| DQA1*04:01+DQB1*04:02 | 416 | 5538 | | DRB1*16:02P | 193 | 5761 | |
| DQA1*05:01+DQB1*02:01 | 1027 | 4927 | | DRB3*01:01P | 1154 | 4800 | DRB3*01:62, DRB3*01:01 |
| DQA1*05:01+DQB1*02:02 | 95 | 5859 | | DRB3*02:02 | 1899 | 4055 | |
| DQA1*05:01+DQB1*03:01 | 183 | 5771 | | DRB3*03:01 | 796 | 5158 | |
| DQA1*05:01+DQB1*05:01 | 129 | 5825 | | DRB4*01:01P | 2370 | 3584 | DRB4*01:01, DRB4*01:03 |
| DQA1*05:01+DQB1*05:03 | 24 | 5930 | | DRB5*01:01 | 1234 | 4720 | |
| DQA1*05:01+DQB1*06:02 | 123 | 5831 | | DRB5*01:02 | 194 | 5760 | |
| DQA1*05:01+DQB1*06:03 | 58 | 5896 | | DRB5*02:02P | 239 | 5715 | DRB5*02:02, DRB5*02:21 |

#### 4 Description of supplementary data

The supplementary data for this manuscript is a zip-compressed tab-separated value file with one row per TCR in the ECOcluster associated with CMV. Its columns have the following interpretations:

- tcr: TCR CDR3 amino acid sequence, V gene and J gene, delimited by '+'
- hla: the HLA allele the TCR is associated with (via its HLA CO-cluster)
- hla_cocluster: the name of the HLA CO-cluster the TCR belongs to
- tcr_pgen: the OLGA generation probability of the TCR
- hla_cocluster_npos_hlamatch: the number of positive-label samples in the holdout CMV-labeled dataset (out of 120) that are inferred to have the associated HLA allele
- hla_cocluster_nneg_hlamatch: the number of negative-label samples in the holdout CMV-labeled dataset (out of 120) that are inferred to have the associated HLA allele
- hla_cocluster_auroc_hlaaware: the AUROC defined by breadth on the HLA-COcluster *vs.* the CMV serological label, considering only donors inferred to have the associated HLA allele
- hla_cocluster_auroc_hlaunaware: the AUROC defined by breadth on the HLA-COcluster *vs.* the CMV serological label, considering all 120 holdout donors
